# Rapid phenotypic and genotypic change in a laboratory schistosome population

**DOI:** 10.1101/2024.08.06.606850

**Authors:** Kathrin S. Jutzeler, Roy N. Platt, Xue Li, Madison Morales, Robbie Diaz, Winka Le Clec’h, Frédéric D. Chevalier, Timothy J.C. Anderson

## Abstract

**Background:** Genomic analysis has revealed extensive contamination among laboratory-maintained microbes including malaria parasites, *Mycobacterium tuberculosis* and *Salmonella* spp. Here, we provide direct evidence for recent contamination of a laboratory schistosome parasite population, and we investigate its genomic consequences. The Brazilian *Schistosoma mansoni* population SmBRE has several distinctive phenotypes, showing poor infectivity, reduced sporocysts number, low levels of cercarial shedding and low virulence in the intermediate snail host, and low worm burden and low fecundity in the vertebrate rodent host. In 2021 we observed a rapid change in SmBRE parasite phenotypes, with a ∼10x increase in cercarial production and ∼4x increase in worm burden.

**Methods:** To determine the underlying genomic cause of these changes, we sequenced pools of SmBRE adults collected during parasite maintenance between 2015 and 2023. We also sequenced another parasite population (SmLE) maintained alongside SmBRE without phenotypic changes.

**Results:** While SmLE allele frequencies remained stable over the eight-year period, we observed sudden changes in allele frequency across the genome in SmBRE between July 2021 and February 2023, consistent with expectations of laboratory contamination. (i) SmLE-specific alleles rose in the SmBRE population from 0 to 41-46% across the genome between September and October 2021, documenting the timing and magnitude of the contamination event. (ii) After contamination, strong selection (*s* = ∼0.23) drove replacement of low fitness SmBRE with high fitness SmLE alleles. (iii) Allele frequency changed rapidly across the whole genome, except for a region on chromosome 4 where SmBRE alleles remained at high frequency.

**Conclusions:** We were able to detect contamination in this case because SmBRE shows distinctive phenotypes. However, this would likely have been missed with phenotypically similar parasites. These results provide a cautionary tale about the importance of tracking the identity of parasite populations, but also showcase a simple approach to monitor changes within populations using molecular profiling of pooled population samples to characterize fixed single nucleotide polymorphisms. We also show that genetic drift results in continuous change even in the absence of contamination, causing parasites maintained in different labs (or sampled from the same lab at different times) to diverge.

## Background

Laboratory research with pathogen populations or cell lines requires rigorous safeguards to prevent contamination and to ensure repeatability of results from different laboratories. Nevertheless, a growing body of literature suggests that contamination (or mislabeling) of laboratory pathogens is surprisingly common. For example, phylogenetic studies of laboratory adapted malaria parasite lines reveal widespread evidence for these issues [1–3]. Contamination from positive control samples have resulted in extensive false positive diagnoses in hospital diagnostic laboratories working with *Mycobacterium tuberculosis*, *Salmonella* spp. and enterococci [4–7]. Finally, methods like isozyme analysis, HLA identity testing, and DNA fingerprinting have exposed misidentification of lymphoma, hematopoietic, and ovarian carcinoma cell lines as a result of cross-contamination [8–10]. In many cases, the contamination may go unnoticed, particularly when no change is observed in pathogen phenotypes or when changes are subtle. As a result, the National Institutes for Health (NIH) and other funding agencies now require provision of protocols for validating the identity of the pathogens under study.

A second process – rapid evolution – can also result in genomic and phenotypic change in pathogen populations over a short time period [11]. Rapid evolution of microbial populations in response to drug pressure, or to avoid immune attack, is ubiquitous. Evolution can also be surprisingly rapid in helminth parasites such as schistosomes. For example, selection for drug resistance [12,13] or cercarial shedding number [14] can substantially alter parasite phenotypes in <10 generations.

The lifecycle of the schistosome parasites can be maintained in the laboratory using freshwater snail intermediate hosts and rodents as definitive hosts. Our laboratory maintains several populations of *Schistosoma mansoni* including two parasite populations originating from Brazil, SmLE and SmBRE.

We have previously investigated these two populations in great detail, and we have reported striking differences in virulence, sporocyst growth, cercarial shedding, and immunopathology between them [15–18]. SmBRE exhibited lower fitness than SmLE for multiple life history traits in both the intermediate and definitive host. However, we noticed a drastic change in phenotypes typical for the SmBRE population starting in 2021. Over time, we noticed increased snail infectivity, higher cercarial shedding, and increased worm burden in SmBRE, while SmLE phenotypes remained relatively unchanged. These observations led us to speculate that the changes observed in the low fitness SmBRE parasites could have resulted from two processes: (i) laboratory contamination with the more efficient SmLE population or (ii) selection of *de novo* mutations within the SmBRE population leading to increased fitness.

To evaluate these alternative scenarios, we sequenced pools of male and female worms from SmBRE and SmLE parasites collected at 10 time intervals over a seven-year period (2016-2023). We monitored allele frequency changes across the genome over time, both within and between the SmBRE and SmLE populations, to answer the following questions: (i) how stable are allele frequencies in laboratory schistosome populations? (ii) Do phenotypic changes in SmBRE reflect selection of *de novo* mutations or laboratory contamination? (iii) If contamination occurred, what can we learn about the dynamics of genomic changes following admixture? (iv) Can we develop molecular approaches to verify laboratory schistosome populations and detect contamination?

## Methods

### Ethics statement

This study was performed in accordance with the Guide for the Care and Use of Laboratory Animals of the National Institutes of Health. The protocol was approved by the Institutional Animal Care and Use Committee of Texas Biomedical Research Institute (permit number: 1419-MA).

### Parasite lifecycle maintenance and recovery of *Schistosoma mansoni* worms

The *S. mansoni* lifecycle spans approximately 75 days (30 days development within snails and 45 days in hamsters). To safeguard against the loss of parasite populations, we establish duplicate cohorts of hamster infections ∼3-4 weeks apart. Many of the same shedding snails are used to infect the two cohorts of hamsters. Hence, the parallel populations of each line form a single population, and some of our adult worm pools are collected one month apart.

To recover adult worms, we perfused infected Golden Syrian hamsters used for schistosome life cycle maintenance as previously described [19]. Briefly, we euthanized each hamster with a solution of 1 ml of phenobarbital (Fatal Plus) + 10% heparin and dissected the animal to expose the liver. We disrupted the hepatic portal vein using a needle and perfused the heart and the liver for around 1 minute each with a perfusion solution (193 nM of NaCl / 1mM EDTA) at a flow rate of 40 ml/minute using a peristaltic pump. After perfusion, we rinsed the intestine with normal saline and collected worms trapped in the intestine. All expelled worms were collected in a fine mesh sieve and rinsed with normal saline solution (154 nM of NaCl, pH 7.5). We then transferred the collected *S. mansoni* worms to a petri dish for counting and separation by sex. The worms were stored in 1.5 ml microcentrifuge tubes, flash-frozen in liquid nitrogen, and preserved at −80 °C until gDNA extraction.

### Cercarial shedding

We used datasets from Le Clec’h et al. [17] from 2015 and performed a similar infection experiment to measure cercarial production of SmBRE in 2023. Briefly, we exposed 240 BgBRE snails to a single SmBRE miracidium in 24-well plates overnight. We then transferred the exposed snails to trays for 4 weeks. At four weeks post-exposure, each snail was individually placed in a well of a 24 well-plate in 1 mL freshwater and kept under artificial light for 2 h to induce cercarial shedding. For each well with cercariae, we sampled three 10 µL (for the high shedder parasites) or 100 µL (for the low shedder parasites) aliquots and added 20 µl of 20× normal saline. We then counted the immobilized cercariae in triplicate under a microscope. We multiplied the mean of the triplicated measurement by the dilution factor to determine the number of cercariae produced by each infected snail. We monitored cercarial production weekly from week 4 to 7 post-exposure in SmBRE-infected snails. To track cercarial production of individual snails throughout the 4-week patent period, we isolated each infected snail in a uniquely labeled 100 mL glass beaker filled with ∼50 mL freshwater at the first shedding. All snails were fed *ad libitum* with fresh lettuce and kept in the dark in the 26-28°C temperature-controlled room.

### gDNA extraction, gDNA Library preparation and sequencing

We extracted gDNA from 27 to 100 single-sex worms per pool (Table 1) with the DNeasy Blood & Tissue Kit (Qiagen, Germantown, MD, USA), following the manufacturer protocol. We ground the worms in 180 μl of ATL buffer using a sterile micro pestle and added 20 μl of proteinase K before incubation at 56°C for 2h. gDNA was eluted in 75 µL of elution buffer. We quantified extracted gDNA using Qubit dsDNA BR Assay Kit (Invitrogen, Carlsbad, CA, USA) and performed library preparation using the KAPA Hyperplus Kit (Roche, Indianapolis, IN, USA) with 400 ng of input material. We used the manufacturer’s instructions with the following modifications for our library construction: enzymatic fragmentation time: 20 minutes, library amplification: six PCR cycles, library size selection: a first size cut at 0.6X (30 µl beads), and a second size cut at 0.8X (10 µl beads). The library sizes were assessed using TapeStation 4200 D1000 ScreenTape (Agilent, Santa Clara, CA, USA), and all libraries were quantified using the KAPA Library Quantification Kit (Roche, Indianapolis, IN, USA). Pooled libraries were submitted to Admera Health and sequenced to high read depth on a NovaSeq X Plus platform (Illumina) with 150 bp paired-end reads.

**Table 1:** Sample information.

| SampleID | Population | Collection Date | Pool size | Sex | Mean coverage | Coverage > 10X | Coverage < 1X | Accession<sup>1</sup> |
| --- | --- | --- | --- | --- | --- | --- | --- | --- |
| BRE110916_m | SmBRE | 11/09/16 | 64 | males | 49.5 | 96.3% | 1.9% | SAMN40565564 |
| BRE110916_f | SmBRE | 11/09/16 | 71 | females | 52.3 | 96.7% | 2.3% | SAMN40565565 |
| LE110216_m | SmLE | 11/02/16 | 77 | males | 48.8 | 96.4% | 1.9% | SAMN40565566 |
| LE110216_f | SmLE | 11/02/16 | 67 | females | 55.4 | 97.8% | 1.4% | SAMN40565567 |
| BRE062718_m | SmBRE | 06/27/18 | 98 | males | 69.4 | 96.8% | 2.0% | SAMN40565568 |
| BRE062718_f | SmBRE | 06/27/18 | 78 | females | 57.3 | 97.6% | 1.5% | SAMN40565569 |
| LE062118_m | SmLE | 06/21/18 | 86 | males | 61.5 | 96.4% | 2.2% | SAMN40565570 |
| LE062118_f | SmLE | 06/21/18 | 87 | females | 50.0 | 96.6% | 2.7% | SAMN40565571 |
| BRE051420_m | SmBRE | 5/14/2020 | 95 | males | 62.8 | 96.2% | 1.7% | SAMN40565572 |
| BRE051420_f | SmBRE | 5/14/2020 | 76 | females | 88.0 | 97.8% | 1.2% | SAMN40565573 |
| LE051420_m | SmLE | 5/14/2020 | 32 | males | 61.6 | 96.6% | 1.3% | SAMN40565574 |
| LE051420_f | SmLE | 5/14/2020 | 30 | females | 51.6 | 97.5% | 1.6% | SAMN40565575 |
| BRE112320_m | SmBRE | 11/23/2020 | 103 | males | 60.0 | 96.0% | 2.1% | SAMN40565576 |
| BRE112320_f | SmBRE | 11/23/2020 | 90 | females | 56.7 | 95.7% | 3.3% | SAMN40565577 |
| LE112320_m | SmLE | 11/23/2020 | 68 | males | 187.7 | 97.3% | 1.2% | SAMN40565578 |
| LE112320_f | SmLE | 11/23/2020 | 106 | females | 56.3 | 97.4% | 1.8% | SAMN40565579 |
| BRE070521_m | SmBRE | 7/5/2021 | 100 | males | 65.7 | 96.4% | 2.2% | SAMN40565580 |
| BRE070521_f | SmBRE | 7/5/2021 | 57 | females | 62.9 | 97.6% | 1.3% | SAMN40565581 |
| LE070521_m | SmLE | 7/5/2021 | 75 | males | 53.7 | 95.2% | 3.1% | SAMN40565582 |
| LE070521_f | SmLE | 7/5/2021 | 73 | females | 54.1 | 96.7% | 2.5% | SAMN40565583 |
| BRE122121_m | SmBRE | 12/21/2021 | 101 | males | 66.9 | 97.7% | 1.5% | SAMN40565584 |
| BRE122121_f | SmBRE | 12/21/2021 | 83 | females | 53.3 | 98.3% | 1.1% | SAMN40565585 |
| LE122121_m | SmLE | 12/21/2021 | 101 | males | 83.9 | 97.5% | 1.8% | SAMN40565586 |
| LE122121_f | SmLE | 12/21/2021 | 104 | females | 68.1 | 97.9% | 1.3% | SAMN40565587 |
| BRE070522_m | SmBRE | 7/5/2022 | 93 | males | 76.2 | 97.6% | 1.8% | SAMN40565588 |
| LE070522_m | SmLE | 7/5/2022 | 64 | males | 69.4 | 96.6% | 1.9% | SAMN40565589 |
| LE070522_f | SmLE | 7/5/2022 | 51 | females | 61.8 | 97.0% | 2.2% | SAMN40565590 |
| BRE021523_m | SmBRE | 2/15/2023 | 96 | males | 93.3 | 97.5% | 1.7% | SAMN40565591 |
| BRE021523_f | SmBRE | 2/15/2023 | 96 | females | 54.6 | 97.6% | 1.9% | SAMN40565592 |
| LE021523_m | SmLE | 2/15/2023 | 106 | males | 66.8 | 97.7% | 1.7% | SAMN40565593 |
| LE021523_f | SmLE | 2/15/2023 | 33 | females | 55.9 | 94.5% | 4.8% | SAMN40565594 |
| BRE092921_m | SmbRE | 9/29/2021 | 100 | males | 90.8 | 96.8% | 2.1% | SAMN40565595 |
| BRE092921_f | SmbRE | 9/29/2021 | 94 | females | 66.1 | 96.8% | 2.1% | SAMN40565596 |
| LE092921_m | SmLE | 9/29/2021 | 27 | males | 71.1 | 96.3% | 1.7% | SAMN40565597 |
| LE092921_f | SmLE | 9/29/2021 | 100 | females | 66.5 | 97.6% | 1.6% | SAMN40565598 |
| BRE102621_m | SmbRE | 10/26/2021 | 100 | males | 79.5 | 97.2% | 1.0% | SAMN40565599 |
| BRE102621_f | SmbRE | 10/26/2021 | 54 | females | 98.0 | 98.5% | 1.0% | SAMN40565600 |
| LE102621_m | SmLE | 10/26/2021 | 60 | males | 93.7 | 97.0% | 1.9% | SAMN40565601 |

<sup>1</sup>All accession numbers are in bioproject PRJNA1090435

### Computational environment

We used conda v23.1.0 to manage environments and download packages required for the analysis. Data processing was performed in R 4.2.0 using tidyverse v1.3.2, and figures were generated with *ggplot* v3.4.2.

### Genotyping

We used trim_galore v0.6.7 [20] (-q 28--illumina--max_n 1--clip_R1 7--clip_R2 7) for adapter and quality trimming before mapping the sequences to version 10 of the *S. mansoni* reference genome (Wellcome Sanger Institute, BioProject PRJEA36577) with BWA v0.7.17-r118 [21] and the default parameters. We used GATK v4.3.0.0 [22] for further processing of the sequences. First, we removed all optical/PCR duplicates with MarkDuplicates. Next, we used HaplotypeCaller and GenotypeGVCFs to call single nucleotide variants (SNV) on a contig-by-contig basis. These were aggregated for each pooled sample and further consolidated into a comprehensive VCF file encompassing all sequences. Quality filtering was performed using VariantFiltration with recommended parameters (FS > 60.0, SOR > 3.0, MQ < 40.0, MQRankSum < −12.5, ReadPosRankSum < −8.0, QD < 2.0). Additionally, we used VCFtools v0.1.16 [23] for refining, specifically excluding non-biallelic sites with quality < 15 and read depth < 10, along with sites and individuals with a genotyping rate < 50%.

We measured selection coefficient (*s*) at each SNP locus by fitting a linear model between the natural log of the allele ratio (freq[allele1]/freq[allele2]) against generation time (measured as the number of 75-day parasite life cycles). The raw *s* values were smoothed by computing the running medians to remove noise.

### F_ST_ statistics

We calculated F_ST_ with popoolation2 [24], a pipeline designed for analysis of pooled samples. Briefly, we used samtools v1.9 [25] mpileup to generate a joint bam file containing sequences from two different samples to make comparisons across time or between populations. Next, we converted the file to a suitable input file for popoolation2 with *mpileup2sync.jar,* keeping only bases with a minimum quality of 20. Finally, we calculated F_ST_ with *fst-sliding.pl* and the following parameters: “--suppress-noninformative”, “--min-count 6”, “--min-coverage 50”, “--max-coverage 200”, “--min-covered-fraction 1”, “--window-size 1”, “--step-size 1”, and the relevant pool sizes with “--pool-size.” We then calculated mean F_ST_ in 20 kb windows using a custom function in R and added the smoothing line using the locfit method from the locfit v1.5-9.8 package.

To calculate F_ST_ for SmLE specific variants, we modified the parameters above to “--min-coverage 10” and “--max-coverage 6000” and overlapped the resulting files with known variant loci.

### Statistical analysis

We performed all statistical analyses with the rstatix v0.7.2 package [26]. For normally distributed data (Shapiro test, *p* > 0.05), we performed parametric Student’s *t*-test to compare time points. Otherwise, we used non-parametric Wilcoxon rank-sum tests. We adjusted *p*-values for multiple comparisons using the Benjamini–Hochberg method when needed and considered these significant when *p* < 0.05 [27].

## Results

### Phenotypic differences between SmBRE parasites from 2015 and 2023

Starting in 2021, we observed an increase in cercarial shedding from infected snails and in worm burden from infected hamsters within the SmBRE population during lifecycle maintenance. As we had previously characterized different SmBRE life history traits, including cercarial shedding in 2015 [17], we repeated this experiment with SmBRE parasites collected in 2023 and quantified cercarial shedding in snails 4-7 weeks post-infection. SmBRE parasites produced 5-17x more cercariae in 2023 than the ones from 2015 (Figure 1A; Week 4: W = 512, *p* < 0.001; Week 5: W = 16.5, *p* < 0.001; Week 6: W = 68, *p* < 0.001; Week 7: W = 9.5, *p* < 0.001).

**Figure 1:**
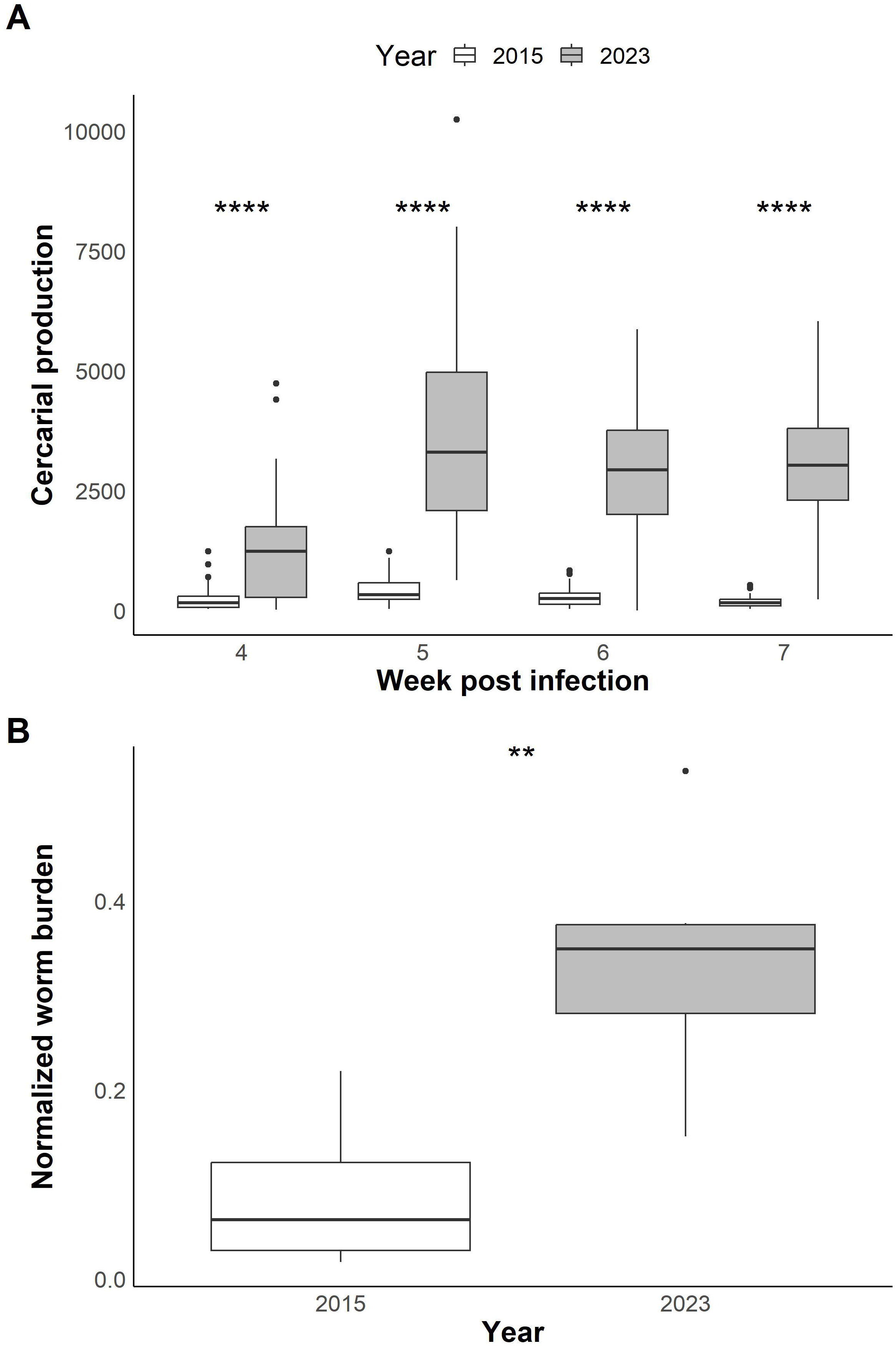
Phenotypic differences between SmBRE and SmLE. **(A)** Boxplots showing cercarial shedding from infected snails, measured in 2015 (data from Le Clec’h et al., 2019) and 2023 over four weeks of the patent period (4-7 weeks post snail infection). Statistical comparisons done between years using a Wilcoxon rank sum test and adjusted for multiple comparisons (Benjamini-Hochberg). **(B)** Boxplots showing worm burden normalized by the number of cercariae used for hamster infection in 2015 and 2023. Statistical comparison between years done with Student’s *t*-test. * *P* < 0.05, ** *P* < 0.01, *** *P* < 0.001, **** *P* < 0.0001.

We used our life cycle maintenance records to quantify changes in worm burden in SmBRE infected hamsters in 2015 and 2023. We normalized worm burden by accounting for variation in the number of cercariae used for hamster infections. We collected almost four times more worms from SmBRE infected hamsters in 2023 compared to their 2015 counterparts (Figure 1A; t_(7.89)_ = −3.55, *p* = 0.008).

### Differentiation between SmBRE and SmLE over time

We used F_ST_ to measure the differentiation between SmBRE and SmLE over time. Genetic markers showed consistent high differentiation (average F_ST_ = 0.24) across the autosomes (chr 1-7) and sex chromosome (chr) Z between 2016 and September 2021 (Figure 2). We observed a drastic reduction in genetic differentiation (F_ST_ reduced from 0.27 to 0.11) between September and October 2021. After October 2021, there was a progressive genome wide reduction in F_ST_ reaching 0.03 by the last sampling date (February 2023).

**Figure 2:**
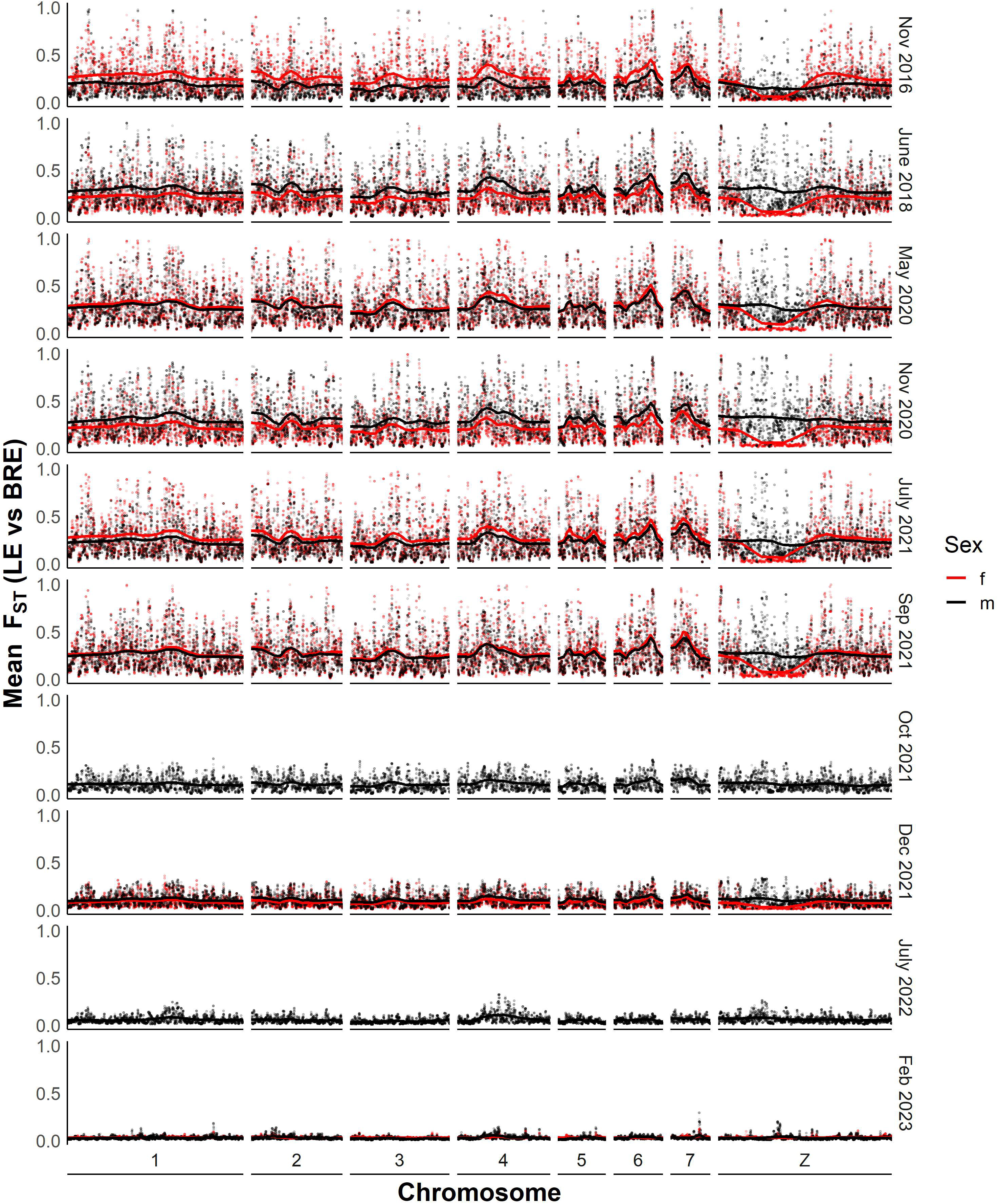
Differentiation between SmBRE and SmLE across time between 2016 and 2023. Dot plot showing smoothed average F_ST_ across the whole genome calculated in 20 kb windows. The solid lines indicate F_ST_ after smoothing with a local regression model as calculated by the locfit R package.

To determine whether SmBRE or SmLE populations were changing over time, we calculated F_ST_ between the earliest time point sampled (2016) and pooled samples from each time point for both SmBRE and SmLE. This information is plotted across the genome in Additional File 1: Figures S1 and Additional File 2: Figure S2 and summarized in Figure 3. This analysis indicates a unidirectional change, stemming from the contamination of SmBRE with SmLE. Across the genome, SmLE parasites showed minor differentiation, with average F_ST_ rising from 0.014 in 2016 to 0.022 in 2023. Meanwhile, we observed a rapid change in SmBRE occurring between September and October 2021, when average F_ST_ suddenly surged from 0.014 to 0.079. From this point on, differentiation intensified, reaching 0.167 by 2023. This significant shift occurred over two years, equivalent to approximately nine 75-day parasite generations.

**Figure 3:**
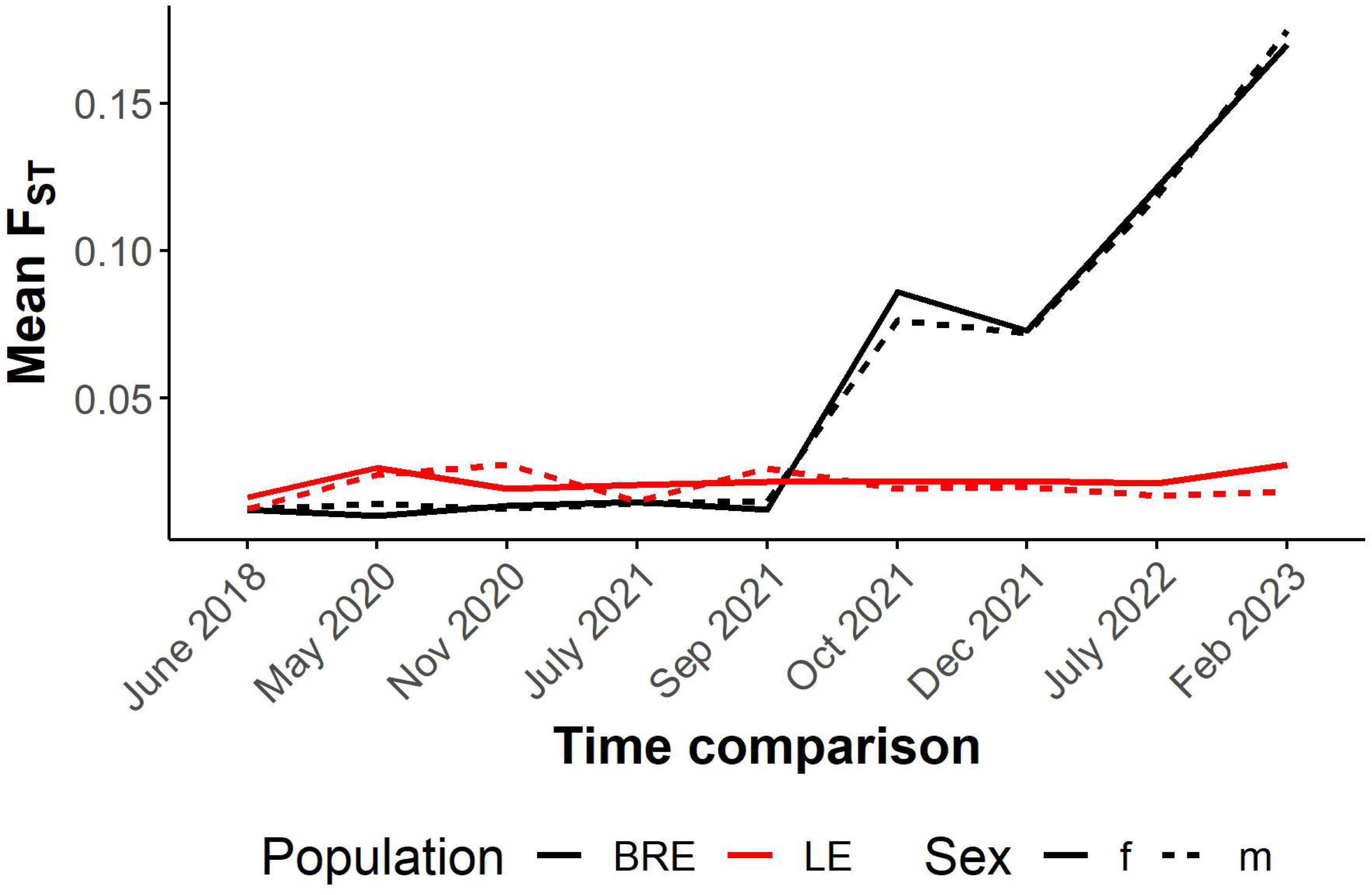
Differentiation in SmBRE and SmLE across time in comparison to 2016. Line plot showing average F_ST_ across the genome for each time point in comparison to pools sampled in 2016.

### Rapid allele frequency change across the SmBRE genome

To more precisely examine the dynamics of this contamination event, we identified 96,778 ancestry informative SNPs that were present in SmLE pools at a frequency of 100% but completely absent in SmBRE pools until September 2021. We then plotted the mean allele frequencies of these SmLE specific loci in all sequenced SmBRE pools (Figure 4A). We saw a consistent jump in mean allele frequency of SmLE specific alleles on each autosome (chr 1-7) and the Z sex chromosome from 0 to 41-46% between September and October 2021, pinpointing when the contamination event occurred and revealing the size of the initial contamination event.

**Figure 4:**
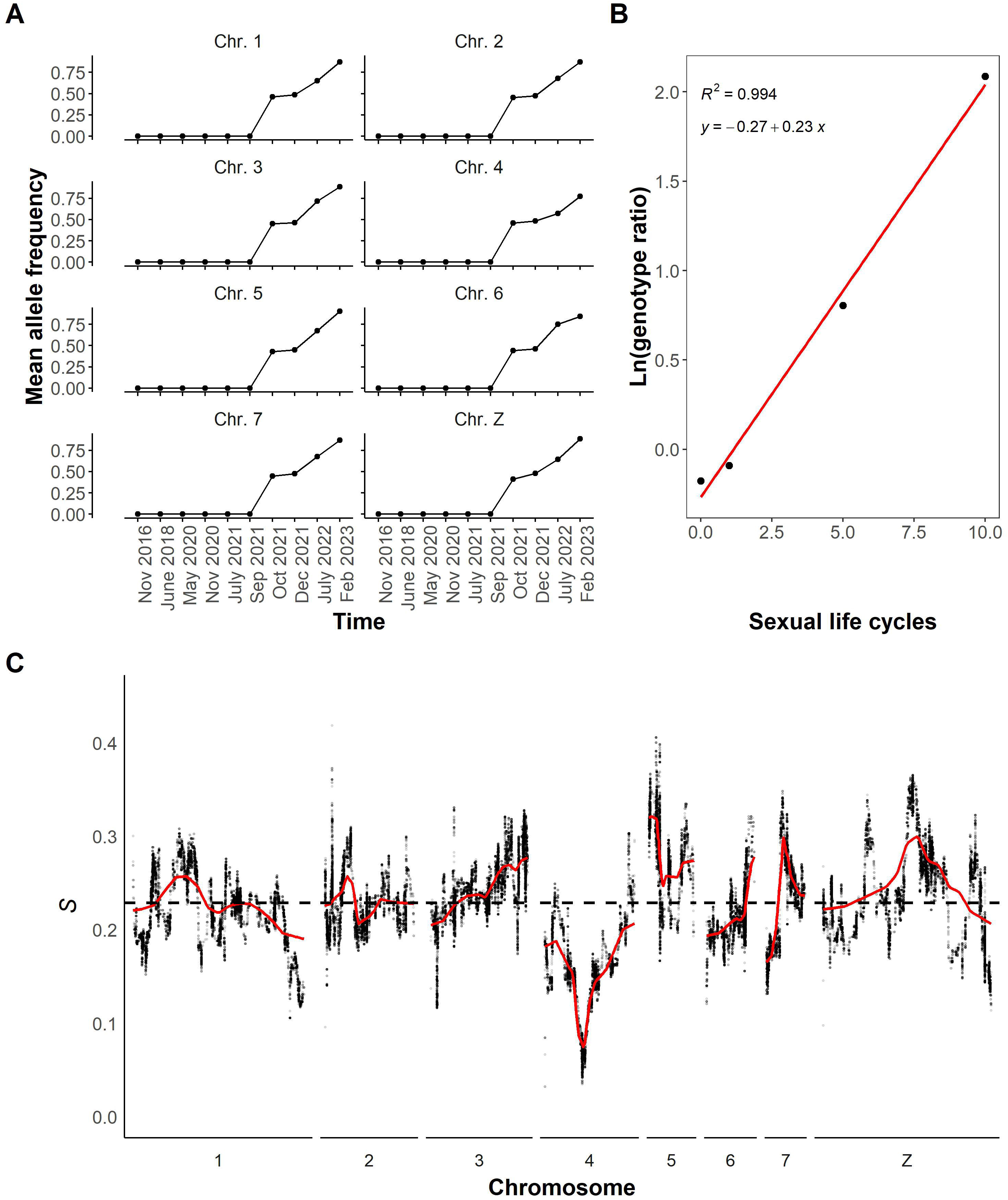
SmLE-specific allele frequencies in SmBRE pools. **(A)** Line plot showing mean allele frequency of SmLE specific variants per chromosome and across time. **(B)** Natural log of the genotype ratio plotted against sexual life cycles. The selection coefficient was estimated as the slope of the least-squares fit. The genotype ratio was calculated as the average genome wide frequency of SmBRE alleles/average genome wide frequency of SmLE alleles at each time point after the initial contamination event. **(C)** Selection coefficient (*s*) for individual SNPs across the whole genome. A local regression smooth line is shown in red.

We also identified 217,657 SmBRE specific variants that were at fixation in SmBRE and absent from SmLE prior to September 2021. These remained undetected in SmLE after September 2021, demonstrating that contamination was unidirectional from SmLE to SmBRE. A summary of SmLE and SmBRE specific SNPs is shown in Table 2, and detailed information for each SNP is listed in Additional File 3: Tables S1 (SmBRE) and Additional File: Table S2 (SmLE).

**Table 2:** Summary of SmBRE and SmLE Specific SNVs.

| Chromosome | Count SmbRE | Count SmLE |
| --- | --- | --- |
| 1 | 43,670 | 16,403 |
| 2 | 32,175 | 11,041 |
| 3 | 21,569 | 14,968 |
| 4 | 24,752 | 8,741 |
| 5 | 19,219 | 8,205 |
| 6 | 13,028 | 14,019 |
| 7 | 12,242 | 7,240 |
| Z | 50,999 | 16,159 |
| MITO | 3 | 2 |
| Total | 217,657 | 96,778 |

### Patterns of selection across the genome

We would expect allele frequencies of SmLE alleles to remain at the same level in subsequent generations, assuming that most introduced SNPs are selectively neutral. However, we observed a steady increase in the frequency of SmLE specific alleles, which reached 77-90% by February 2023. The average patterns of change are extremely similar across the genome (Figure 4A), with the exception of chr 4 where we observed slower change.

To investigate allele frequency change across the genome after the initial contamination event, we calculated selection coefficients (*s*) for SmLE specific SNPs. Figure 4B shows average changes in allele frequency of SmLE across the genome (plotted as the natural log of the genotype ratio) against time (in parasite generations) and reveals a good fit to a linear model, with a slope of 0.23, demonstrating strong selection towards SmLE alleles across the genome. We then calculated selection coefficients for individual SmLE specific SNPs and plotted these across the genome (Figure 4C). Selection coefficients for SmLE specific alleles average *s* = 0.23 across the whole genome as expected, but there are peaks where *s* = 0.41 on chr 5, and *s* = 0.37 on the Z chr. There is a 1.55 Mb region of particular interest on chr 4, where *s* < 0.06, and frequencies of SmBRE alleles showed minimal change following the initial admixture event. This was the only genome region where selection for *SmLE* alleles was weak (*s* between 0.03 and 0.06). This region contains 11 genes (Additional File 5: Table S3).

### Changes in allele frequency in SmLE parasite pools

The seven-year longitudinal series of SmLE samples provides an opportunity to examine stability of allele frequencies over time in the absence of contamination. There were 706,496 SNPs segregating within our SmLE populations. Variant SNPs were defined as those showing genetic variation (MAF > 0.05) in at least one of the time periods sampled. While some of these show large changes in allele frequencies over the 7-year dataset (Figure 5A), the majority remain stable over time, as shown by F_ST_ comparisons of 2016 pools with 2023 pools (Figure 5B). Similarly, allele frequency changed by 0.16 in males and 0.17 in females on average between 2016 and 2023 (Figure 5C). However, 0.31% of segregating SNPs showed allele frequency change of > 0.8, while 0.08% spread to fixation.

**Figure 5:**
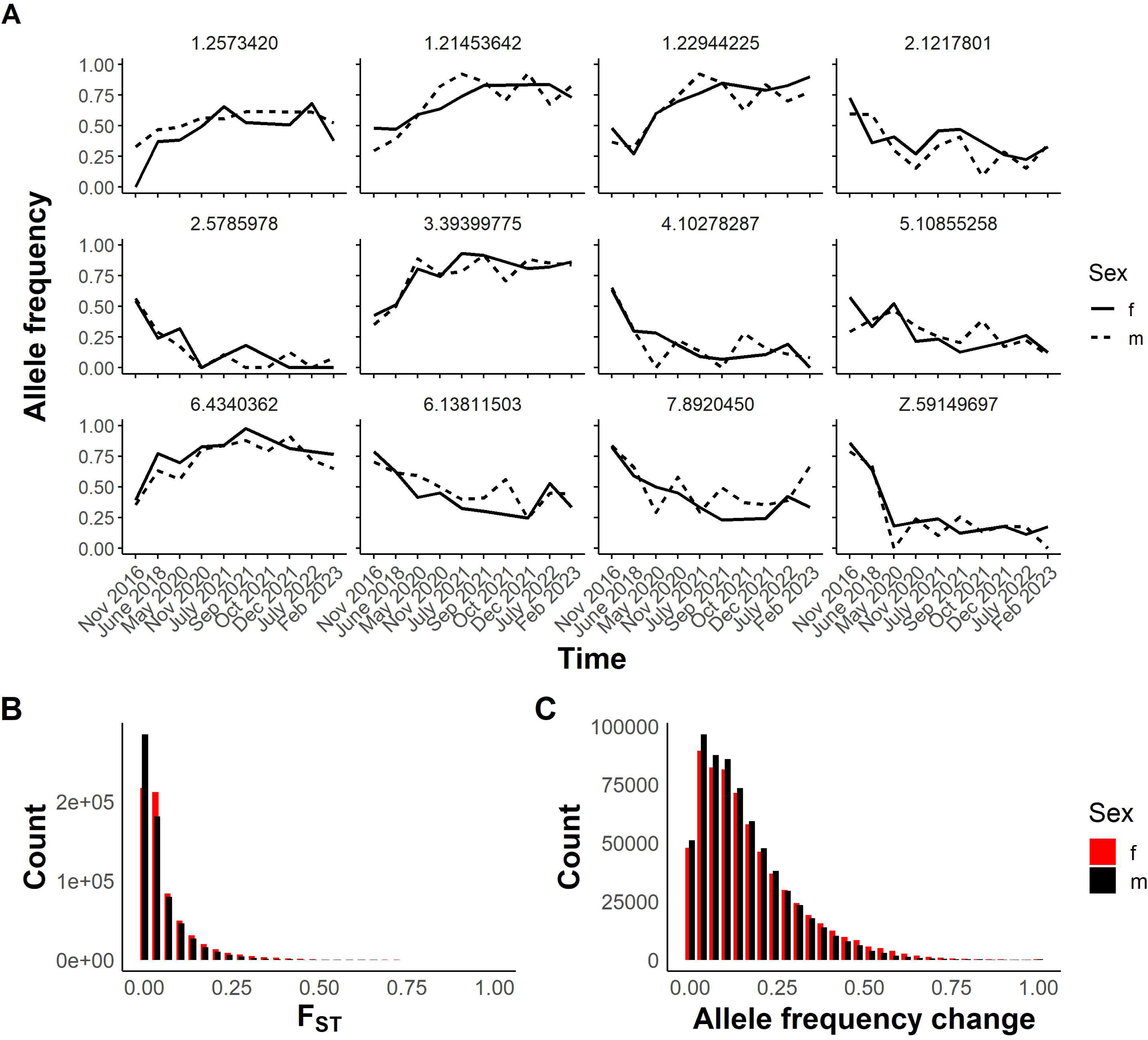
Observed differentiation in SmLE parasites over time. **(A)** Line plot showing allele frequency change over time in specific variants in the SmLE population. Variants are labeled by chromosome and position. **(B)** Histogram illustrating the distribution of F_ST_ values from the comparison of 657,592 variants in female pools and 661,996 variants in male pools from 2016 with those from 2023. **(C)** Distribution of allele frequency change in the same variants between 2016 and 2023.

## Discussion

Using pooled sequencing analyses, we demonstrate that the drastic increase in SmLE-specific alleles in the SmBRE population, resulted from a unidirectional contamination event, with SmLE taking over the SmBRE population except for a singular region on chr 4. We speculate that mixing cercariae or miracidia during life cycle maintenance was the cause of this contamination event, as we performed this task for both populations at the same time.

### Dynamics of a laboratory contamination event

#### Size of initial contamination event

We observed a 40-46% change in the frequency of SmLE specific markers in the SmBRE population in a single generation. The change is of the same magnitude across the autosomes and the Z chr. This indicates that 40-46% of worms analyzed from October 2021 resulted from infection with SmLE rather than SmBRE cercariae. The actual proportion of SmLE cercariae in the infecting pools was likely much lower than 40-46%, because SmLE shows 1.8 fold higher establishment rate than SmBRE [28]. Assuming that contamination occurred during the cercariae stage, we therefore speculate that the contaminating fraction was ∼22.2 – 25.6%.

#### Genome replacement of SmBRE by SmLE alleles

After contamination, we might expect that allele frequencies from each parent would remain relatively stable in the absence of selection. However, our analyses show a systematic genome-wide increase in SmLE specific alleles over time. Selection is extremely strong (*s* = 0.23) averaged across the whole genome. This is comparable to selection for artemisinin resistance in *P. falciparum* [29]. To further put this in perspective, the estimated mean selection coefficient in humans is ∼ 0.001, targeting only 1% of the genome [30]. We have previously determined quantitative trait loci (QTLs) on chr 1, 3, and 5 that underlie high cercarial shedding rates in SmLE; these were identified through genetic crosses with SmBRE [16]. We predicted that these regions would show a rapid increase in SmLE specific alleles, but that other genome regions would remain unchanged. Instead, we see a consistent increase in proportion of SmLE specific SNPs across the genome in the admixed population. The genome-wide changes observed suggest limited mating between SmLE and SmBRE worms within admixed populations. Such assortative mating may occur due to differences in establishment rate of mature worms in the blood vessels. We speculate that SmLE establishes in the portal venous system before SmBRE, and that SmLE males and females are already paired and producing eggs prior to emergence of mature SmBRE adults. As a consequence, SmLE eggs are overrepresented in the liver eggs that are harvested to found the next generation, leading to the genome-wide replacement of SmBRE with SmLE alleles. We note that fecundity is also three times greater in SmLE than SmBRE females [15]. This will further accelerate replacement of SmBRE alleles by SmLE and contributes to the high genome wide selection (*s* = 0.23) for SmLE alleles.

#### Variation in strength of selection across the genome

Some regions of the genome show higher or lower selection coefficients than the genome wide average of 0.23. This suggests that some mating between SmBRE and SmLE occurs, and that some genome regions show much stronger selection. Of particular interest is the region on chr 4. This is the only part of the genome in which SmBRE alleles remain at high frequency after the initial admixture event (Table S3). The chr 4 region contains 11 genes, including a Leishmanolysin-like peptidase (Smp_127030). This class of metalloprotease-encoding genes impact infection rates in both snail and vertebrate host [31,32].

We also observe four genome regions (chr 2, 5, 7, and Z) showing particularly high selection coefficients indicating extremely strong selection for SmLE alleles (Table S3). These regions do not correspond to the QTLs determining cercarial production in previous genetic crosses between SmBRE and SmLE [16].

### Genetic drift in SmLE parasites

We saw no evidence for contamination in SmLE. The seven-year longitudinal data set from this population provides a valuable opportunity to examine allele frequency change due to genetic drift. We observed a subset of SNPs present in SmLE pools exhibiting high allele frequency changes over the seven-year period. We have previously determined that laboratory schistosome populations retain abundant genetic variation (Jutzeler et al., unpublished observations). In the SmLE parasite pools examined here, there are 706,496 SNPs with allele frequency > 5%. SNPs changed in allele frequency on average by 0.16 between 2016 and 2023. However, variance was high and a subset (0.31%) of segregating SNPs changed in frequency by > 0.8 between 2016-2023. The effective population size *N_e_* in laboratory maintained *S. mansoni* populations is relatively small (53-264 in SmLE, Jutzeler et al. unpublished observations). While some of the change in allele frequencies may be driven by selection, the pattern observed is broadly consistent with genetic drift and results in gradual change in the SmLE population over the years. These results illustrate how parasite populations maintained in different laboratories, or sampled from the same laboratory over time, may differ in allele frequency. Hence, the reproducibility of experiments may potentially be affected simply by the divergence of the schistosome populations.

### Pooled sequencing for validating schistosome populations and identifying contamination

Developing a simple approach to characterize laboratory schistosome populations is challenging because these populations show abundant genetic variation (Jutzeler et al., unpublished observations). Sequencing pools of parasites provides a simple and relatively inexpensive solution, because we can profile SNVs that are fixed within populations. These SNVs should remain relatively stable indicators of population identity baring contamination, or mutation, which is expected to be extremely rare. Here, we share a list of population specific SNPs (Table S1 and S2) to help with the identification and validation of SmBRE and SmLE parasite populations. Expanding these lists to include other commonly used schistosome parasite populations would provide an important resource for verifying the identity of these populations and detecting potential contamination.

### Implications for schistosome research

How commonly does contamination occur in laboratory schistosome populations? In addition to the event documented in this paper, we have also retrospectively discovered a contamination of the SmHR parasite population, which was fixed for the *SmSULT-OR Δ142* mutation responsible for oxamniquine resistance (Winka Le Clec’h and Frederic Chevalier, unpublished observations). We received the SmHR population in 2016 but found that the *SmSULT-OR Δ142* mutation was no longer at 100% frequency, most likely as a result of contamination. We therefore used marker-assisted selection to “purify” this population (now named SmOR) by conducting single miracidium infections and established hamster infections with cercariae that were fixed for the Δ142 deletion. Hence, there are a minimum of two known contamination events in laboratory schistosome populations.

Schistosomes are typically maintained by laboratory passage through its hosts, because cryopreservation, while possible, is quite inefficient [33]. As a result, even if such contamination events occur extremely rarely, they can cause irreversible changes to the genetic makeup of laboratory parasite populations. Moreover, these changes may go undetected if they don’t alter specific phenotypes. The results observed in SmLE, where no contamination occurred, also demonstrate how genetic drift within parasite populations can lead to gradual change in allele frequencies. Characterizing pooled population samples using fixed SNP profiles of pooled parasites, as described here, will be a powerful tool to verify parasite identity and determine the extent of contamination and the magnitude of change resulting from genetic drift in laboratory parasite populations.

How does the contamination event documented here impact interpretation of prior experiments using SmBRE? We recently used SmBRE and other parasite populations, to investigate the contribution of parasite and host genotype on immunopathology in the mouse host [15]. The cercariae used for rodent infections in this experiment were obtained from snails infected in July 2021, prior to the contamination event. Hence, this experiment was unaffected. We also examined genetic variation in five distinct *S. mansoni* populations (Jutzeler et al., unpublished observations). This work was conducted after the contamination event, but we replaced the SmBRE parasites used initially with −80°C-preserved SmBRE worms collected prior to contamination to avoid this issue.

We note that the snail intermediate hosts used for maintaining schistosome populations in the laboratory are also maintained as continuously breeding colonies and cannot currently be cryopreserved. Like schistosomes, these snail colonies are maintained as genetically variable, sexually reproducing populations, and contamination between co-maintained colonies is a potential issue. We suggest that profiles of fixed SNPs could also provide a valuable approach to detecting contamination and maintaining integrity of laboratory snail populations.

### Conclusions

This study demonstrates a significant contamination event between the SmBRE and SmLE parasite populations, leading to a notable increase in SmLE-specific alleles within the SmBRE population. The potential for genetic drift within these populations, as evidenced by the gradual changes in allele frequencies in the SmLE population, further underscores the necessity for tools to validate the identity of laboratory-maintained schistosome populations.

## Supplementary information

**Additional file 1: Figure S1. Differentiation of SmBRE parasites between 2016 and all following time points.** Dot plot showing smoothed average F_ST_ across the whole genome calculated in 20 kb windows. The solid lines indicate F_ST_ after smoothing with a local regression model as calculated by the locfit R package.

**Additional file 2: Figure S2. Differentiation of SmLE parasites between 2016 and all following time points.** Dot plot showing smoothed average F_ST_ across the whole genome calculated in 20 kb windows. The solid lines indicate F_ST_ after smoothing with a local regression model as calculated by the locfit R package.

**Additional file 3: Table S1. List of SmBRE specific variants.** The reference alleles are those shown at each position listed in version 10 of the *S. mansoni* reference genome (Wellcome Sanger Institute, BioProject PRJEA36577).

**Additional file 4: Table S2. List of SmLE specific variants.** The reference alleles are those shown at each position listed in version 10 of the *S. mansoni* reference genome (Wellcome Sanger Institute, BioProject PRJEA36577).

**Additional file 5: Table S3. Genes under selection.** This table lists the genes and corresponding gene ontology (GO) terms as identified by WormBase’s BioMart v0.7 [34].

## Availability of data and materials

The datasets supporting the conclusions of this article and all codes used for data analysis and generation of figures (1-5, S1-S2) are available at https://github.com/kathrinsjutzeler/BRE-LE-contamination and Zenodo 10.5281/zenodo.13136643. Sequencing data is available on NCBI short read archive (SRA), under BioProject PRJNA1090435 (accession numbers: SAMN40565564 to SAMN40565601, Table 1).

## Authors’ contributions

KSJ and TJCA designed and planned the experiments. WL, FDC infected snails and counted cercariae. WL, FDC, MM and RD maintained parasites and collected pools of adult worms. KSJ performed experimental and molecular work and analyzed data. RNP and XL provided guidance on data analysis. KSJ and TJCA drafted the manuscript. All authors read and approved the final manuscript.

## Acknowledgements

We thank Evelien Bunnik, Elizabeth Leadbetter, Robin Leach, and P’ng Loke for insightful comments and suggestions on this work.

## REFERENCES

1. Mu J, Awadalla P, Duan J, McGee KM, Joy DA, McVean GAT, et al. Recombination hotspots and population structure in Plasmodium falciparum. PLoS Biol. 2005;3:e335.

2. Nair S, Nkhoma S, Nosten F, Mayxay M, French N, Whitworth J, et al. Genetic changes during laboratory propagation: copy number At the reticulocyte-binding protein 1 locus of *Plasmodium falciparum*. Mol Biochem Parasitol. 2010;172:145–8.

3. Neafsey DE, Schaffner SF, Volkman SK, Park D, Montgomery P, Milner DA, et al. Genome-wide SNP genotyping highlights the role of natural selection in Plasmodium falciparum population divergence. Genome Biol. 2008;9:R171.

4. Jasmer RM, Roemer M, Hamilton J, Bunter J, Braden CR, Shinnick TM, et al. A prospective, multicenter study of laboratory cross-contamination of Mycobacterium tuberculosis cultures. Emerg Infect Dis. 2002;8:1260–3.

5. De Lappe N, Connor JO, Doran G, Devane G, Cormican M. Role of subtyping in detecting Salmonella cross contamination in the laboratory. BMC Microbiol. 2009;9:155.

6. de Boer AS, Blommerde B, de Haas PEW, Sebek MMGG, Lambregts-van Weezenbeek KSB, Dessens M, et al. False-positive mycobacterium tuberculosis cultures in 44 laboratories in The Netherlands (1993 to 2000): incidence, risk factors, and consequences. J Clin Microbiol. 2002;40:4004–9.

7. Katz KC, McGeer A, Low DE, Willey BM. Laboratory contamination of specimens with quality control strains of vancomycin-resistant enterococci in Ontario. J Clin Microbiol. 2002;40:2686–8.

8. Liscovitch M, Ravid D. A case study in misidentification of cancer cell lines: MCF-7/AdrR cells (re-designated NCI/ADR-RES) are derived from OVCAR-8 human ovarian carcinoma cells. Cancer Lett. 2007;245:350–2.

9. Drexler HG, Dirks WG, MacLeod RA. False human hematopoietic cell lines: cross-contaminations and misinterpretations. Leukemia. 1999;13:1601–7.

10. Drexler HG, MacLeod RA, Dirks WG. Cross-contamination: HS-Sultan is not a myeloma but a Burkitt lymphoma cell line. Blood. 2001;98:3495–6.

11. Messer PW, Petrov DA. Population genomics of rapid adaptation by soft selective sweeps. Trends Ecol Evol. 2013;28:659–69.

12. Couto FFB, Coelho PMZ, Araújo N, Kusel JR, Katz N, Jannotti-Passos LK, et al. *Schistosoma mansoni*: a method for inducing resistance to praziquantel using infected *Biomphalaria glabrata* snails. Mem Inst Oswaldo Cruz. 2011;106:153–7.

13. Rogers SH, Bueding E. Hycanthone resistance: development in Schistosoma mansoni. Science. 1971;172:1057–8.

14. Gower CM, Webster JP. Fitness of indirectly transmitted pathogens: restraint and constraint. Evolution. 2004;58:1178–84.

15. Jutzeler KS, Le Clec’h W, Chevalier FD, Anderson TJC. Contribution of parasite and host genotype to immunopathology of schistosome infections. Parasit Vectors. 2024;17:203.

16. Le Clec’h W, Chevalier FD, McDew-White M, Menon V, Arya G-A, Anderson TJC. Genetic architecture of transmission stage production and virulence in schistosome parasites. Virulence. 2021;12:1508–26.

17. Le Clec’h W, Diaz R, Chevalier F, McDew-White M, Anderson T. Striking differences in virulence, transmission and sporocyst growth dynamics between two schistosome populations. Parasites & Vectors. 2019;12:485.

18. Le Clec’h W, Chevalier FD, Jutzeler K, Anderson TJC. No evidence for schistosome parasite fitness trade-offs in the intermediate and definitive host. Parasites Vectors. 2023;16:132.

19. Tucker MS, Karunaratne LB, Lewis FA, Freitas TC, Liang Y. Schistosomiasis. Current Protocols in Immunology [Internet]. 2013 [cited 2020 Nov 10];103. Available from: https://onlinelibrary.wiley.com/doi/abs/10.1002/0471142735.im1901s103

20. Krueger F, James F, Ewels P, Afyounian E, Weinstein M, Schuster-Boeckler B. TrimGalore [Internet]. Available from: https://github.com/FelixKrueger/TrimGalore

21. Li H, Durbin R. Fast and accurate short read alignment with Burrows–Wheeler transform. Bioinformatics. 2009;25:1754–60.

22. McKenna A, Hanna M, Banks E, Sivachenko A, Cibulskis K, Kernytsky A, et al. The Genome Analysis Toolkit: A MapReduce framework for analyzing next-generation DNA sequencing data. Genome Res. 2010;20:1297–303.

23. Danecek P, Auton A, Abecasis G, Albers CA, Banks E, DePristo MA, et al. The variant call format and VCFtools. Bioinformatics. 2011;27:2156–8.

24. Kofler R, Orozco-terWengel P, De Maio N, Pandey RV, Nolte V, Futschik A, et al. PoPoolation: a toolbox for population genetic analysis of next generation sequencing data from pooled individuals. PLoS One. 2011;6:e15925.

25. Danecek P, Bonfield JK, Liddle J, Marshall J, Ohan V, Pollard MO, et al. Twelve years of SAMtools and BCFtools. GigaScience. 2021;10:giab008.

26. Kassambara A. rstatix: Pipe-Friendly Framework for Basic Statistical Tests [Internet]. 2023. Available from: <https://CRAN.R-project.org/package=rstatix>

27. Benjamini Y, Hochberg Y. Controlling the False Discovery Rate: A Practical and Powerful Approach to Multiple Testing. Journal of the Royal Statistical Society: Series B (Methodological). 1995;57:289– 300.

28. Jutzeler KS, Le Clec’h W, Chevalier FD, Anderson TJC. Contribution of parasite and host genotype to immunopathology of schistosome infections [Internet]. Microbiology; 2024 Jan. Available from: http://biorxiv.org/lookup/doi/10.1101/2024.01.12.574230

29. Li X, Kumar S, McDew-White M, Haile M, Cheeseman IH, Emrich S, et al. Genetic mapping of fitness determinants across the malaria parasite Plasmodium falciparum life cycle. PLoS Genet. 2019;15:e1008453.

30. Zeng J, Xue A, Jiang L, Lloyd-Jones LR, Wu Y, Wang H, et al. Widespread signatures of natural selection across human complex traits and functional genomic categories. Nat Commun. 2021;12:1164.

31. Hambrook JR, Hanington PC. A cercarial invadolysin interferes with the host immune response and facilitates infection establishment of Schistosoma mansoni. PLoS Pathog. 2023;19:e1010884.

32. Hambrook JR, Kaboré AL, Pila EA, Hanington PC. A metalloprotease produced by larval Schistosoma mansoni facilitates infection establishment and maintenance in the snail host by interfering with immune cell function. PLoS Pathog. 2018;14:e1007393.

33. Stirewalt M, Cousin CE, Lewis FA, Leefe JL. Cryopreservation of Schistosomules of *Schistosoma Mansoni* in Quantity *. The American Journal of Tropical Medicine and Hygiene. 1984;33:116–24.

34. Consortium WP. WormBase ParaSite BioMart [Internet]. Available from: https://parasite.wormbase.org/biomart/martview/91ea287e9ed5f190f9da26ae4d9a9ba3

