## Supplementary figures and images for "Rapid phenotypic and genotypic change in a laboratory schistosome population"

### Supplemental Figure 1

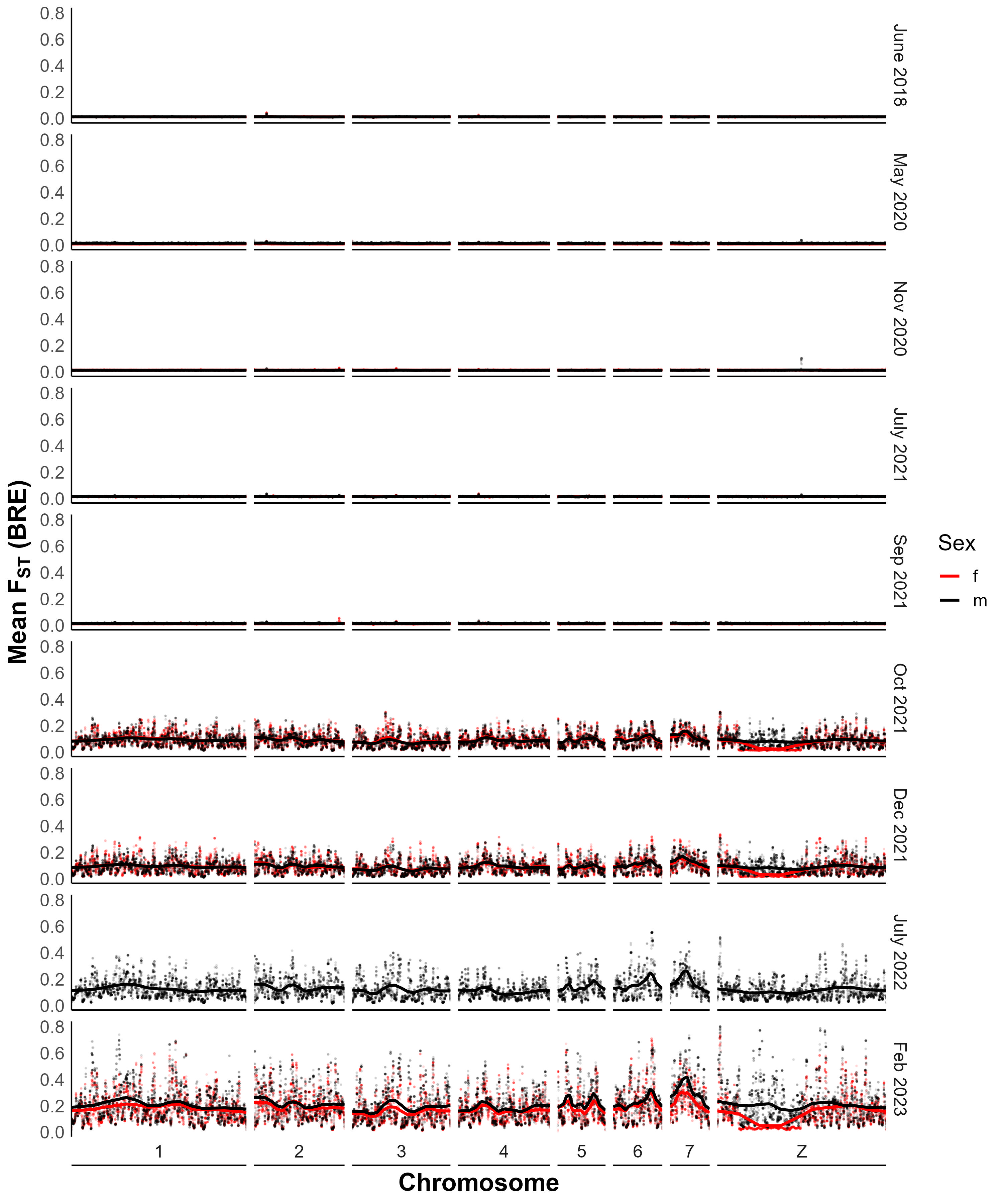

### Supplemental Figure 2

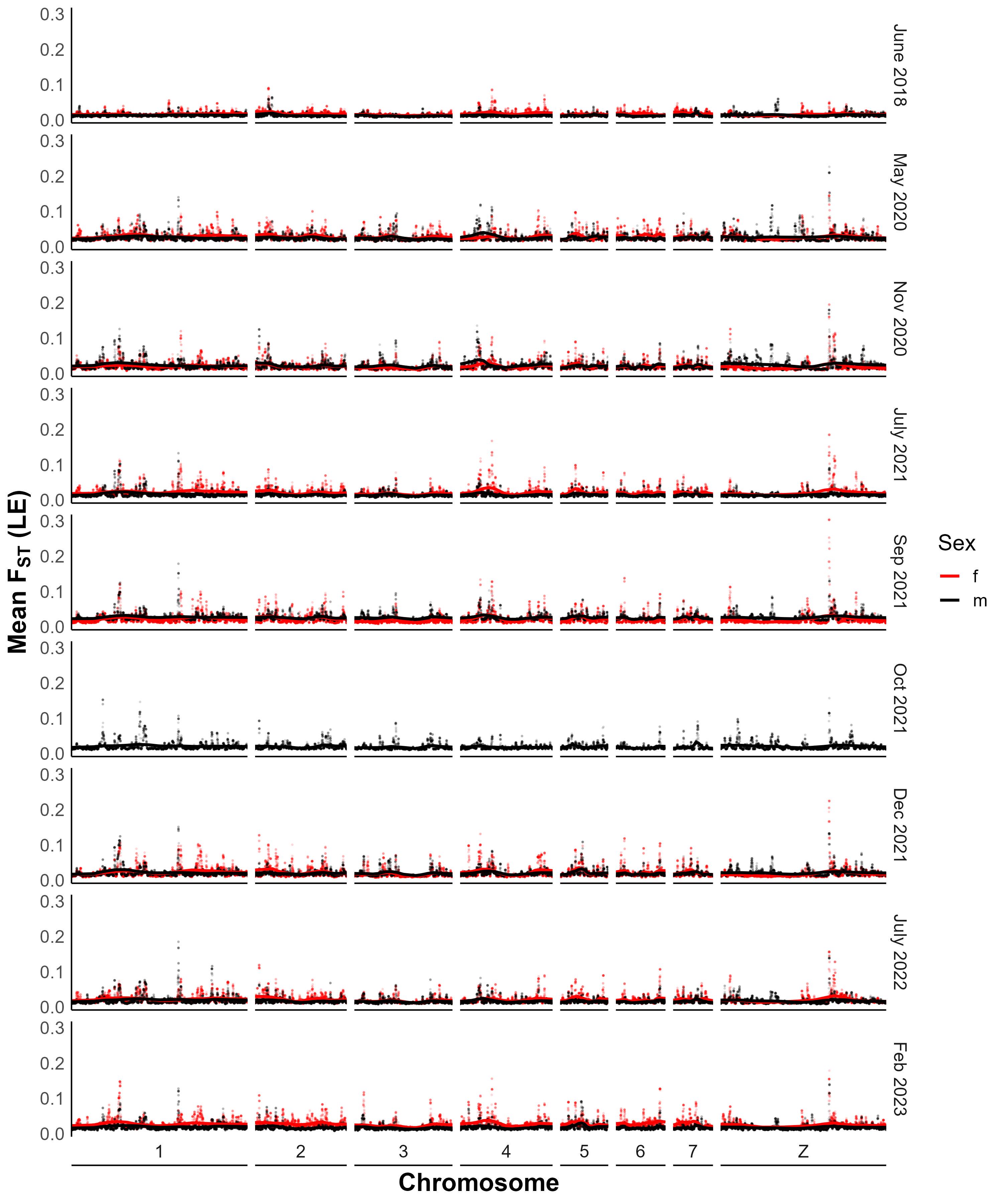
